# Tumor SETDB1 expression predicts worse overall survival for bladder cancer patients

**DOI:** 10.64898/2026.10.23.761403

**Authors:** Sarah Athans, Meegan Young, Han Yu, Bhavisha Doshi, Ellen Karasik, Shuhei Ueshima, Jia Fang, Anna Woloszynska

## Abstract

Aberrant activity of histone methyltransferases promotes tumorigenesis across multiple cancer types, making this class of enzymes attractive therapeutic targets. The histone methyltransferase SETDB1 is frequently amplified in bladder cancer and has been implicated in chemotherapy resistance. We examine whether SETDB1 protein expression is associated with overall survival and tumor characteristics in bladder cancer. To assess SETDB1 protein expression in patients, we performed immunohistochemistry in tissue microarrays (TMAs) containing 350 bladder cancer cases with well-annotated clinical information. Digital image analysis was used to quantify SETDB1 protein expression to produce a SETDB1 H-score, from which patients were categorized as SETDB1-high or SETDB1-low. Using these categories, we compared tumor characteristics, clinical outcomes, and gene expression profiles between SETDB1-high and SETDB1-low patients. Patients with high SETDB1 expression (n=143) had significantly shorter overall survival compared to SETDB1-low (n=125) patients (median survival months: SETDB1-high, 30.09; SETDB1-low, 87.85; p = 0.0006), and SETDB1 expression was independently associated with overall survival upon multivariate cox regression analysis (HR = 1.93, 95% CI = 1.08 - 3.45, p = 0.026). SETDB1-high patients were more likely to present with squamous (35% vs 20%, p = 0.026) and sarcomatoid (11% vs 3.6%, p=0.049) differentiation features, which are associated with poorer prognosis. Exploratory analyses identified transcriptional differences associated with SETDB1 expression, providing rationale for further exploration of the mechanisms by which SETDB1 may promote bladder cancer.

## Introduction

Bladder cancer (BCa) is a highly prevalent disease globally and represents the most common tumor of the urinary tract and the ninth most common cancer worldwide [1]. In 2026, an estimated 84,530 new cases of BCa and 17,870 deaths are projected to occur in the United States, with approximately 77% of the burden observed in males [2]. Although BCa disproportionately affects older adults, its public health burden is projected to rise substantially: projections from WHO demonstrate the absolute number of cases are expected to double by 2050 [3]. BCa is typically treated with cisplatin-based chemotherapy, most commonly a combination of gemcitabine and cisplatin [4]. Because responses to cisplatin-based regimens vary among patients, effort has been made to identify molecular predictors of treatment sensitivity; for example, ATM, RB1, FANCC, or ERCC2 gene mutations have been suggested to predict response to neoadjuvant cisplatin-based chemotherapy [5]. However, clinically useful biomarkers for predicting treatment response and prognosis in BCa remain limited, emphasizing the importance of identifying and developing prognostic biomarkers.

SETDB1 (SET domain bifurcated 1), an H3K9-specific histone methyltransferase, is located on human chromosome 1q21.3 [6]. The SETDB1 protein belongs to the SET domain protein methyltransferase family and mediates H3K9 mono-, di-, and trimethylation in euchromatic regions, thereby regulating gene expression, particularly silencing, through dynamic modulation of chromatin structure [7]. Its physiological role has additionally been associated with de novo epigenetic silencing [8], stem cell maintenance [9], and endogenous retrovirus gene repression [10]. SETDB1 upregulation has been implicated in the tumorigenesis and progression of various human carcinomas, including melanoma, colorectal cancer, ovarian cancer, non-small cell lung cancer (NSCLC) and breast cancer [11–15], wherein high expression was correlated with worse overall survival. SETDB1 also promotes metastasis in cancers such as NSCLC and hepatocellular carcinoma [16, 17]. Furthermore, abnormal expression of SETDB1 has been suggested as a therapeutic target and/or diagnostic biomarker of patient survival in NSCLC [18], and it has been shown that inhibition of SETDB1 can sensitize tumors to immune checkpoint blockade therapy [19], further supporting its emerging relevance as a therapeutic target. Currently, one molecule with specificity for targeting SETDB1 has been discovered: SETDB1- TTD-IN-1 (also known as (R,R)-59) [20]. However, this molecule binds to the tandem tudor domains within the SETDB1 protein instead of the catalytic SET domain, and thus relevance of this compound to cancer treatment may be limited.

While SETDB1 has been increasingly implicated in tumor progression and treatment response across multiple malignancies, its expression patterns and clinical significance in BCa warrant further investigation. Previous work demonstrated that SETDB1 promotes invasion and migration of bladder cancer cells *in vitro* and facilitates EMT-mediated BCa metastasis and gemcitabine resistance [21]. Further investigation of its expression in an expanded cohort of patient tumor samples is warranted to further assess SETDB1 associated tumor characteristics and quantify the prognostic value of SETDB1 expression in bladder cancer. The present study quantified SETDB1 expression in BCa using SETDB1-stained tissue microarrays (TMAs) from 350 BCa patients, thereby providing additional insight into its potential role in bladder tumor biology. Improved understanding of SETDB1 expression patterns may help clarify its relevance to BCa initiation, progression, and response to therapy, and provide rationale for further development of SETDB1-specific inhibitors.

## Materials and Methods

Five tissue microarrays (TMAs) were obtained with informed consent from 377 BCa patients treated at Roswell Park Comprehensive Cancer Center in Buffalo, NY. Patient characteristics can be found in Table 1. Of these, 350 cases had sufficient clinical information to proceed with analyses. Of the 350 cases with clinical information, 269 had sufficient tumor material within the associated microarray cores for immunohistochemical quantification of SETDB1.

**Table 1.** Characteristics of the tissue microarray cohort. All evaluable characteristics for the patients included in the tissue microarray cohort (n = 350). Data presented as median (Q1, Q3) or n (%).

| Characteristic | Total N = 350 |
| --- | --- |
| SETDB1 H-Score | 12 (1, 54) |
| Unknown | 82 |
| AGE_DX | 69 (62, 76) |
| Unknown | 1 |
| SEX |  |
| FEMALE | 79 (23%) |
| MALE | 270 (77%) |
| Unknown | 1 |
| RACE |  |
| AMERICAN INDIAN ALEUT ESKIMO | 1 (0.3%) |
| BLACK | 9 (2.6%) |
| WHITE | 339 (97%) |
| Unknown | 1 |
| ETHNICITY |  |
| NON-SPANISH | 332 (100%) |
| SPANISH, NOS; HISPANIC, NOS; LATINO, NOS | 1 (0.3%) |
| Unknown | 17 |
| ALCOHOL_HIST |  |
| CURRENT USE OF ALCOHOL | 96 (47%) |
| NO HISTORY OF ALCOHOL USE | 95 (46%) |
| PAST HISTORY OF ALCOHOL USE | 15 (7.3%) |
| Unknown | 144 |
| TOBACCO_HIST |  |
| CURRENT | 104 (30%) |
| NEVER USED | 61 (18%) |
| PREVIOUS USE | 182 (52%) |
| Unknown | 3 |
| PATH_AJCC_STAGE |  |
| 0 | 65 (19%) |
| 1 | 40 (11%) |
| 2 | 44 (13%) |
| 3 | 76 (22%) |
| 4 | 124 (36%) |
| Unknown | 1 |
| Squamous_Differentiation | 59 (25%) |
| Unknown | 117 |
| Glandular_Differentiation | 19 (8.2%) |
| Unknown | 117 |
| Sarcomatoid_Features_Differentiation | 15 (6.4%) |
| Unknown | 117 |
| Neuroendocrine_Differentiation | 2 (0.9%) |
| Unknown | 130 |
| Micropapillary_Differentiation | 8 (3.6%) |
| Unknown | 130 |
| Bladder_Cancer_Registry_Cancer_Recurrence_GRIN2A | 27 (8.2%) |
| Unknown | 22 |
| Chr6_24_Mb |  |
| Amplified | 59 (18%) |
| Not amplified | 264 (82%) |
| Unknown | 27 |
| Amplified_cases_GRIN2A |  |
| Amplified | 28 (8.6%) |
| Not amplified | 299 (91%) |
| Unknown | 23 |
| HER2 |  |
| Amplified | 24 (7.6%) |
| Not amplified | 290 (92%) |
| Unknown | 36 |
| chr5_40mb |  |
| Amplified | 34 (10%) |
| Not amplified | 303 (90%) |
| Unknown | 13 |
| FGFR1_amplified |  |
| Amplified | 18 (5.3%) |
| Deleted | 1 (0.3%) |
| Not amplified | 318 (94%) |
| Unknown | 13 |
| chr_remodel_muts | 10 (38%) |
| Unknown | 324 |
| HMT_muts | 13 (50%) |
| Unknown | 324 |
| HAT_muts | 6 (23%) |
| Unknown | 324 |
| Number_Amplified |  |
| 0 | 218 (67%) |
| 1 | 70 (21%) |
| 2 | 27 (8.3%) |
| 3 | 10 (3.1%) |
| 4 | 2 (0.6%) |
| Unknown | 23 |
| Chr5_or_6_Amplified |  |
| 1 | 68 (86%) |
| 2 | 11 (14%) |
| Unknown | 271 |
| Chr6_duplicates |  |
| Yes | 16 (100%) |
| Unknown | 334 |

### Tissue Microarray Staining for SETDB1 by Immunohistochemistry

The TMAs were stained for SETDB1 at a dilution of 1:500 using SETDB1 antibodies derived from rabbit using purified antigens [900–1152aa (C)] and affinity purified from serum [16]. Tissues were fixed in 10% buffered formalin for 24 hours before processing. Tissues were embedded in paraffin and sectioned at 5 microns. Slides were de-paraffinized in several baths of xylene and then rehydrated in graded alcohols followed by ddH_2_O. Using DAKO PT Link module, slides were incubated in 1x pH6 citrate buffer (Invitrogen Cat #00-5000) for 20 minutes. Slides were then incubated in 3% H2O2 for 15 min in DAKO Autostainer. To block non-specific binding, tissues were incubated with 10% normal goat serum for 10 min, followed by avidin/biotin block (Vector Labs Cat#SP-2001). Primary antibody SETDB1 (1:500) was diluted in Envision Flex antibody diluent (Agilent #DM830) and incubated for 30 minutes at room temperature, followed by the biotinylated Goat anti-Rabbit secondary antibody (Vector Biolabs #BA-1000) for 15 minutes. For signal enhancement, ABC reagent (Vector Labs Cat #PK 6100) was applied for 30 minutes. To reveal endogenous peroxidase activity, slides were incubated with DAB substrate (Dako Cat #K3467) for 5 minutes and then counterstained with DAKO Hematoxylin for 20 seconds. Slides were dehydrated through several baths of graded alcohols and xylenes and then coverslipped.

The stained slides were then Aperio scanned by Pathology Core Facility. Each TMA was visually analyzed with ImageScope Positive Pixel Count v9 Algorithm. Areas of tumor cells were manually annotated for each case, including triplicates and excluding controls. Tumor identification was based on histological criteria for anaplasia, such as pleomorphism, hyperchromasia, nuclear enlargement, loss of polarity, increased and/or atypical mitoses, multinucleation, and invasion. All quantified annotations were saved in ImageScope files for visual review or future reanalysis.

### SETDB1 Expression Quantification

3,3’-diaminobenzidine (DAB) nuclear staining for SETDB1 was quantified using a calibrated digital image analysis algorithm using Aperio ImageScope (Leica Biosystems). Individual tumor cell nuclei were identified, and staining intensity was measured following color deconvolution to separate the DAB chromogen from the hematoxylin counterstain. Based on predefined intensity thresholds, nuclei were classified into four categories: 0 (>220; negative), 1+ (220; Weak), 2+ (195; Moderate), or 3+ (175; Strong). For each region of interest, the algorithm determines the number and percentage of cells within each staining category (0, 1+, 2+ and 3+), the percentage of positively stained cells, and the H-Score. The H-score ranges from 0 to 300 using the formula: 1 x (%1+) + 2 x (%2+) + 3 x (%3+). A score of 300 indicates that 100% of tumor cells exhibit strong (3+) nuclear staining. Hematoxylin served as the counterstain to provide tissue morphology and facilitate accurate identification of nuclear structures during image analysis.

### Biostatistical Analysis and Correlation with Clinical Outcomes

A total of 350 patients were included in this analysis. Baseline characteristics are summarized as median with interquartile range (IQR) for continuous variables and frequency with percentage for categorical variables. H-scores were unavailable for 82 patients due to a lack of detectable tumor cells in the TMA core. Because the H-score exhibited a skewed distribution, it was transformed using log_2_ 1(x + 1) for visualization and regression analyses. One patient had unknown survival data and was therefore excluded from survival analyses.

For categorical analyses, the H-score was dichotomized using an optimal cutoff of 10 determined by the maximally selected log-rank statistic with respect to overall survival (OS). Overall survival was estimated using the Kaplan–Meier method, and survival distributions were compared using the log-rank test. Associations between H-score, clinical covariates, and OS were first evaluated using univariable Cox proportional hazards regression. Characteristics with a univariable p-value < 0.10 were considered for inclusion in the multivariable Cox proportional hazards model, which included age at diagnosis, alcohol history, pathologic American Joint Committee on Cancer (AJCC) stage, glandular differentiation, chromosome 6 24 Mb amplification, FGFR1 amplification, and number of detected amplifications. Race and ethnicity were excluded from the regression analyses because of limited variability within the study cohort.

### SETDB1 Predictive Performance

The predictive performance of SETDB1 for overall survival (OS) was evaluated using the concordance index (C-index), which quantifies the model’s ability to correctly rank patients according to their survival risk. Higher C-index values indicate better discriminative ability. To further assess the biomarker’s time-dependent predictive performance, time-dependent receiver operating characteristic (ROC) curve analyses were performed at 2, 3 and 5 years. The corresponding areas under the ROC curves (AUCs) were calculated to quantify the biomarker’s ability to discriminate between patients who experienced the event and those who remained event-free at each time point.

### RNA Sequencing, Differential Expression Analysis, and Gene Set Enrichment Analysis

Single-end raw sequencing reads were first pre-processed by using fastqc (v0.10.1)[22] and mapped to RefSeq GRCh38 human reference genome and corresponding gene annotation obtained from UCSC’s repository using splicing-detection tools Bowtie [23] and TopHat (v2.0.13)[24]. A second pass QC was done using alignment output with RSeQC [25] to examine abundances of genomic features, splicing junction saturation and gene-body coverage. Gene expression is quantified using HTSeq [26] using the -m intersection-strict option. Differential expression analyses were performed using DESeq2 (v1.18.1)[27] to detect differential gene expression in RNA-seq data based on SETDB1 classification with a pre-determined H-score cut-off of 10. Age at diagnosis was included as a clinical covariate for differential expression analysis. Downstream analyses and visualization plots were done using regularized-log2 transformation implemented by DESeq2. Enrichment analysis was performed using gene set enrichment analysis (GSEA; v4.1.0) for Hallmark and curated gene sets, with a minimum of 1,000 permutations per dataset. Permutations were based on gene set.

## Results

### Elevated SETDB1 expression is associated with shorter overall survival for bladder cancer patients

SETDB1 is frequently amplified and/or upregulated at the mRNA level in publicly available clinical bladder cancer datasets [28, 29] (Figure S1A), suggesting that SETDB1 may contribute to bladder cancer initiation or progression. However, whether this corresponds with alterations in patient survival remains unknown. Therefore, we sought to assess protein expression of SETDB1 in bladder cancer clinical specimens. We performed immunohistochemistry staining for SETDB1 in a tissue microarray (TMA) bladder cancer cohort containing cases from 350 patients, then correlated SETDB1 expression level with patient and tumor characteristics. Of the 350 patients, 269 had sufficient tumor cell content on the TMA for downstream analyses based on SETDB1. Patient characteristics can be found in Table 1. Tumor cells in three cores per patient sample were manually circled, and tumor cells were scored based on strength of SETDB1 immunohistochemistry (IHC) staining using a nuclear quantification algorithm. SETDB1 histo-scores (H-scores) were calculated by averaging the expression scores for all evaluable tumor cores per patient, resulting in an H-score between zero (no expression in evaluated tumor cells) to 300 (strong expression in all evaluated tumor cells). Tumor core SETDB1 expression H-scores ranged from 0 to 217.715 and demonstrate a left-skewed distribution (Figure 1). Representative images demonstrating tumor cores with high and low nuclear SETDB1 expression are shown in Figure S1B.

**Figure 1.**
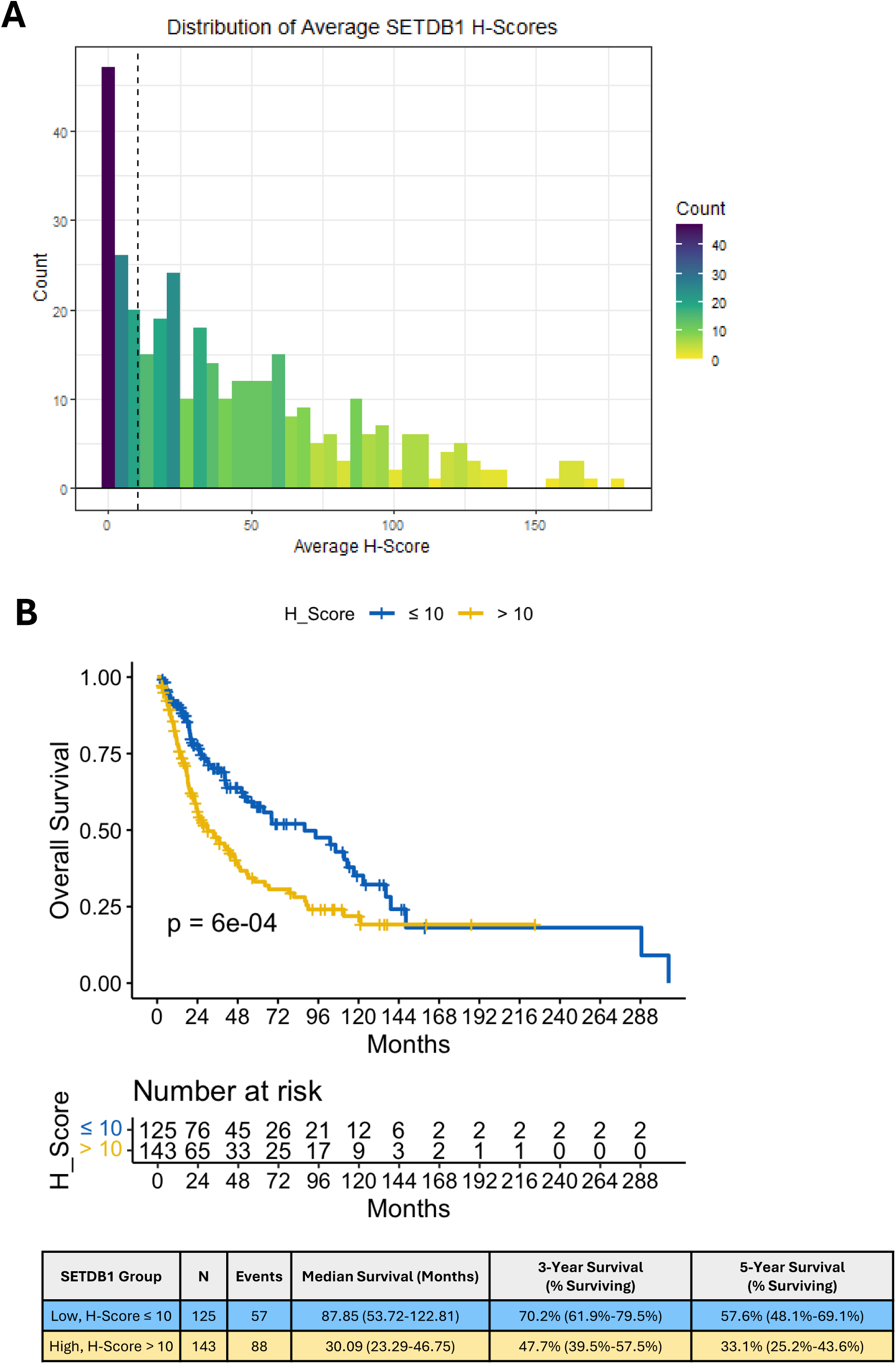
High SETDB1 expression associated with shorter survival. **A.** Histogram showing the frequency of average SETDB1 H-score for the evaluated tumor cores in the tissue microarray. **B.** Top: Kaplan-Meier overall survival curve for patients with high SETDB1 protein expression (H-score > 10; n = 143) and low SETDB1 protein expression (H-score ≤ 10; n = 125). Risk table shows the number of patients at risk in each group at each indicated time point in months. P-value determined by log-rank test. Bottom: Table of median survival months, percentage of survival at 3 years, and percentage of survival at 5 years survival for patients in the SETDB1 high and low groups. Parentheses indicate 95% confidence intervals for each metric in each group.

To determine the optimal H-score cutoff for stratifying patient survival, we evaluated the calculated log-rank test statistics and selected the maximum value, which corresponds to the cutoff with the largest separation in survival curve between above- and below-cutoff groups. From this, we determined an H-score of 10 to be the optimal cutoff to stratify patient survival outcomes (Figure S1C). Based on an H-score of 10, we stratified patients with available survival data (n = 268) into SETDB1-high (n=143) and SETDB1-low (n=125) groups, then compared overall survival and found that SETDB1-high patients have significantly shorter survival compared to the SETDB1-low group (median survival months, SETDB1-high: 30.09; SETDB1- low: 87.85, p = 6*10^-4^, Figure 1B). In addition to SETDB1 expression, we found that age at diagnosis (HR = 1.02, 95% CI =1.01-1.04), American Joint Committee on Cancer (AJCC) tumor stage of 2 or greater (Stage 2 HR (95% CI): 1.95 (1.04, 3.67), Stage 3: 3.57 (2.10, 6.09), Stage 4: 4.84 (2.98, 7.87), past history of alcohol use (HR = 2.16, 95% CI = 1.07, 4.36), and absence of chromosome 6 24Mb amplification (HR = 0.67, 95% CI = 0.46, 0.98) were significantly associated with altered survival outcomes (Table 2). To control for possible confounding factors, we further evaluated the association of SETDB1 expression with overall survival using multivariable cox regression analysis. SETDB1 group (>10 vs ≤10) H-scores and clinical variables with a p-value < 0.10 in the univariable analyses were included as covariates in the multivariable Cox proportional hazards models. Upon multivariable analysis, SETDB1 expression remained significantly associated with shorter patient survival (HR = 1.93, 95% CI = 1.08 - 3.45, p = 0.026, Table 3). This demonstrates a strong significant association between SETDB1 and overall survival for bladder cancer patients, even when controlling for other clinically relevant variables.

**Table 2.**
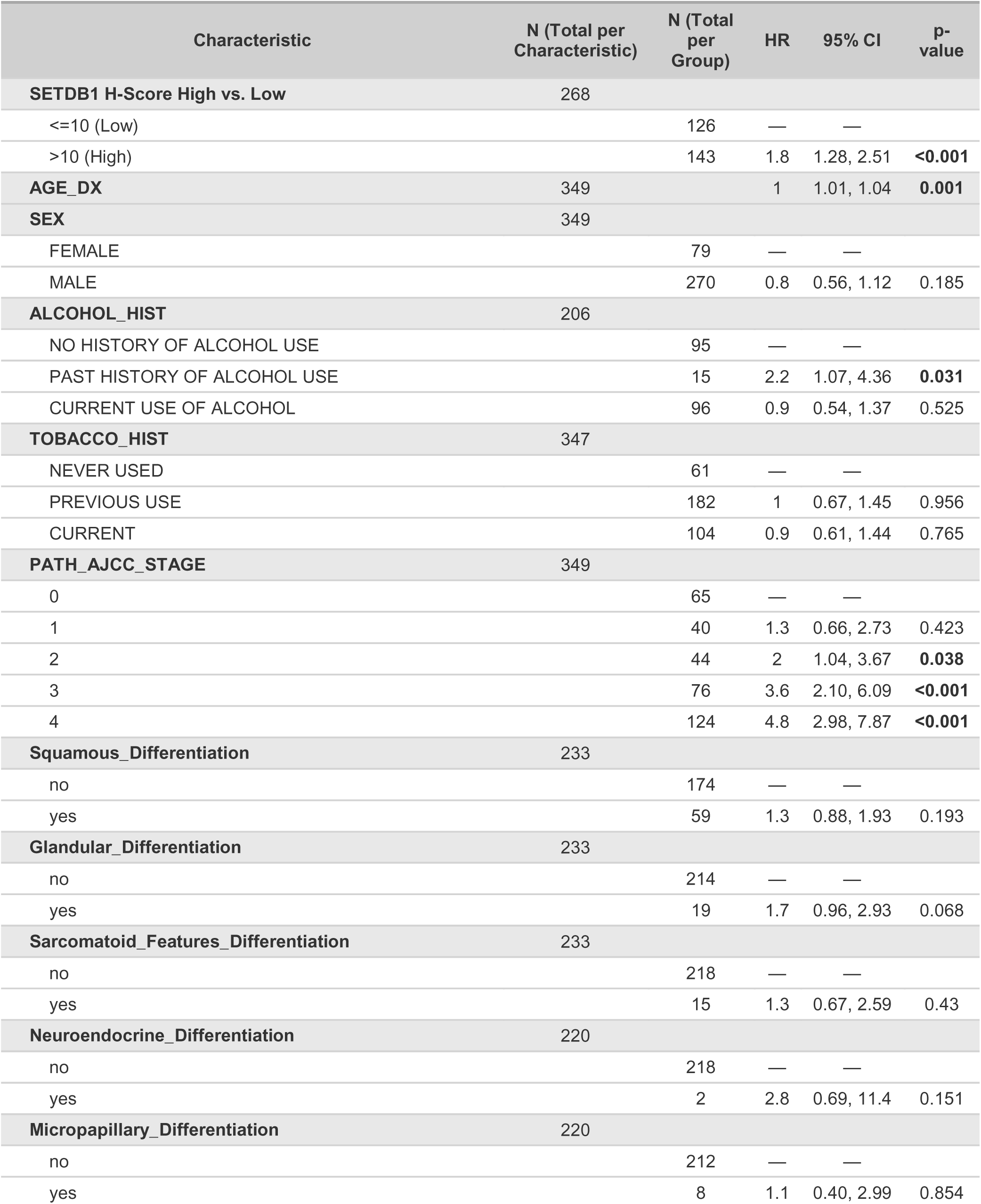
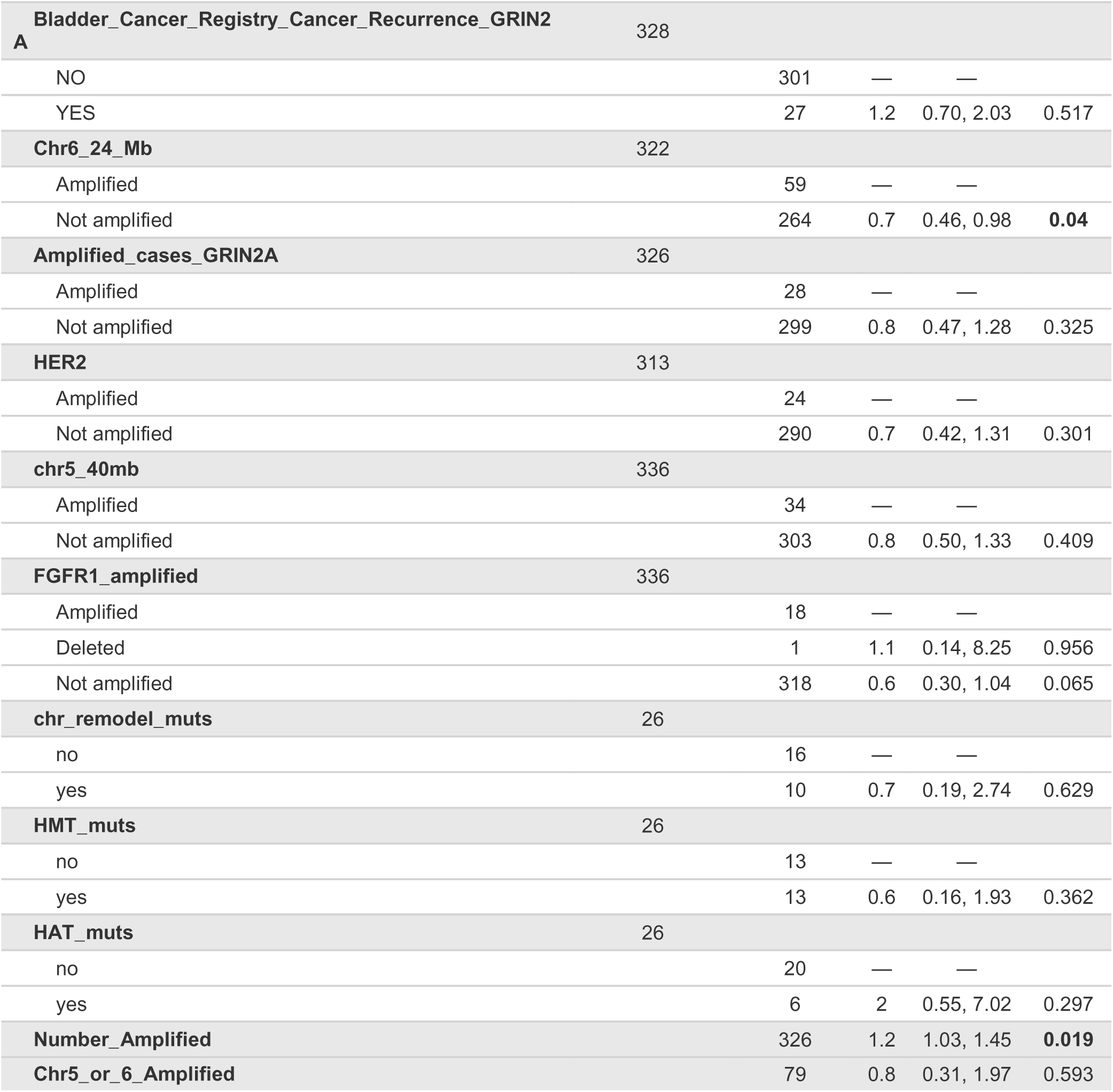
Univariable cox regression analysis of overall survival with clinical variables. Results of the univariable Cox proportional hazards analysis evaluating the associations between overall survival (OS) and H-score as well as other clinical variables (n=268 patients). Abbreviations: CI = Confidence Interval, HR = Hazard Ratio.

**Table 3.** Multivariable cox regression analysis of overall survival. Results of multivariable Cox proportional hazards analyses including a categorical H-score with a cutoff of 10 (n=247 patients). Abbreviations: CI = Confidence Interval, HR = Hazard Ratio.

| Characteristic | HR | 95% CI | p-value |
| --- | --- | --- | --- |
| **SETDB1 H-Score High vs. Low** |  |  |  |
| <=10 | — | — |  |
| >10 | 1.93 | 1.08, 3.45 | **0.026** |
| **AGE_DX** | 1.02 | 1.01, 1.04 | 0.009 |
| **ALCOHOL_HIST** |  |  |  |
| NO HISTORY OF ALCOHOL USE | — | — |  |
| PAST HISTORY OF ALCOHOL USE | 2.52 | 1.21, 5.28 | **0.014** |
| CURRENT USE OF ALCOHOL | 0.78 | 0.45, 1.34 | 0.371 |
| Unknown | 0.98 | 0.62, 1.56 | 0.935 |
| **PATH_AJCC_STAGE_2** |  |  |  |
| <=2 | — | — |  |
| >2 | 2.38 | 1.39, 4.07 | **0.002** |
| **Glandular_Differentiation** |  |  |  |
| no | — | — |  |
| yes | 1.58 | 0.83, 3.00 | 0.165 |
| Unknown | 0.51 | 0.28, 0.93 | **0.029** |
| **Chr6_24_Mb** |  |  |  |
| Amplified | — | — |  |
| Not amplified | 1.19 | 0.65, 2.15 | 0.575 |
| **FGFR1_amplified** |  |  |  |
| Amplified | — | — |  |
| Deleted | 3.3 | 0.35, 31.0 | 0.295 |
| Not amplified | 0.51 | 0.23, 1.12 | 0.094 |
| **Number_Amplified** | 0.92 | 0.66, 1.29 | 0.644 |

To determine the predictive potential of SETDB1 on patient survival, we performed Receiver Operating Characteristic (ROC) analysis, analyzing the predictive power at 2, 3, and 5-year intervals. We found that SETDB1 expression modestly predicts survival at all evaluated time points, with the strongest effect seen at 2 years (Figure S1D).

### SETDB1 expression is associated with patient and tumor characteristics

To assess whether SETDB1 expression was associated with differences in patient or tumor characteristics, we compared the distribution of SETDB1-high and SETDB1-low cases across available clinical variables. Age, sex, race, ethnicity, and alcohol history distributions were not different between SETDB1 low and high groups (Table 4). Tobacco use history differed between the SETDB1-high and SETDB1-low patients (p=0.02; Table 4).

**Table 4.**
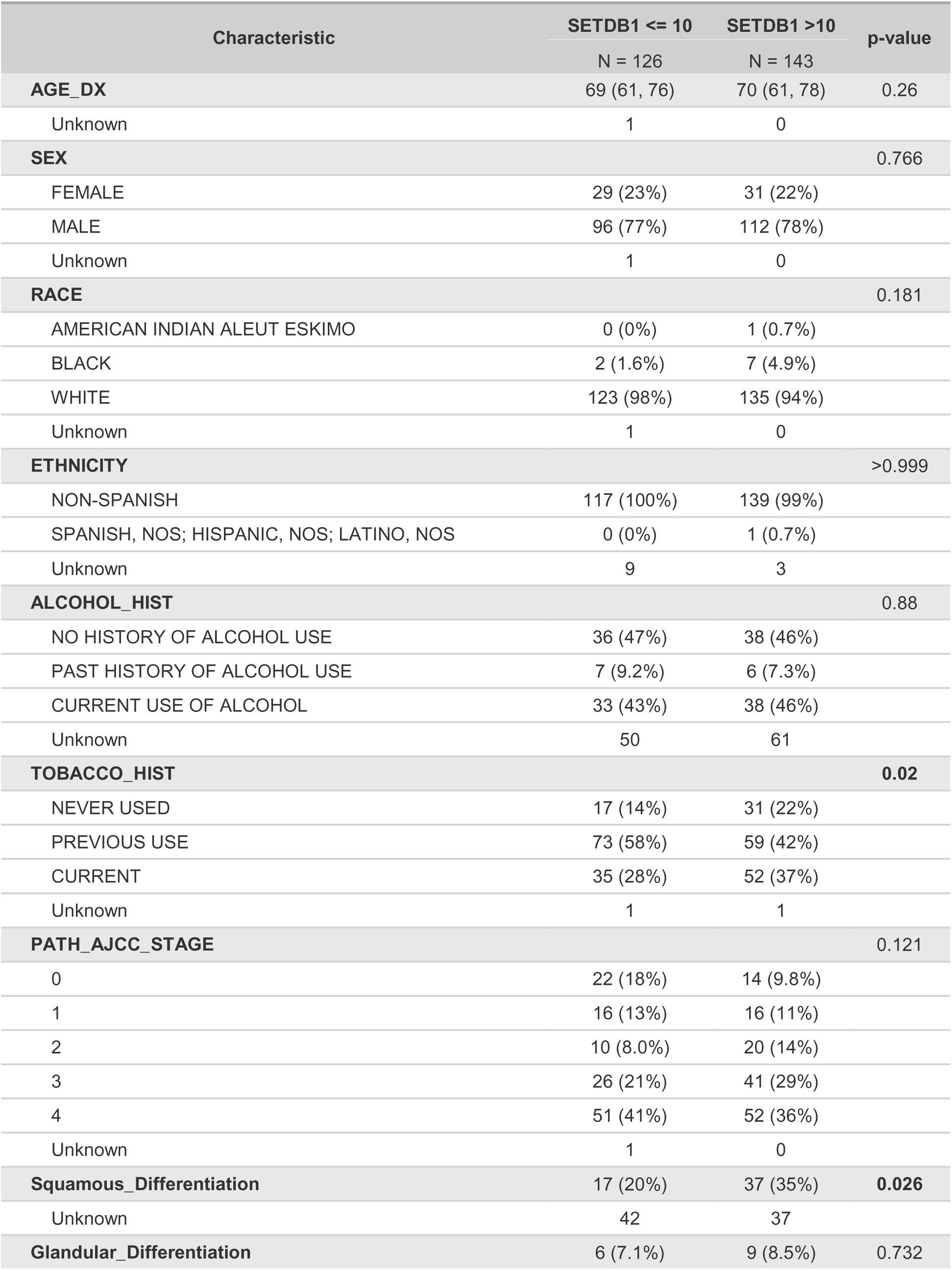
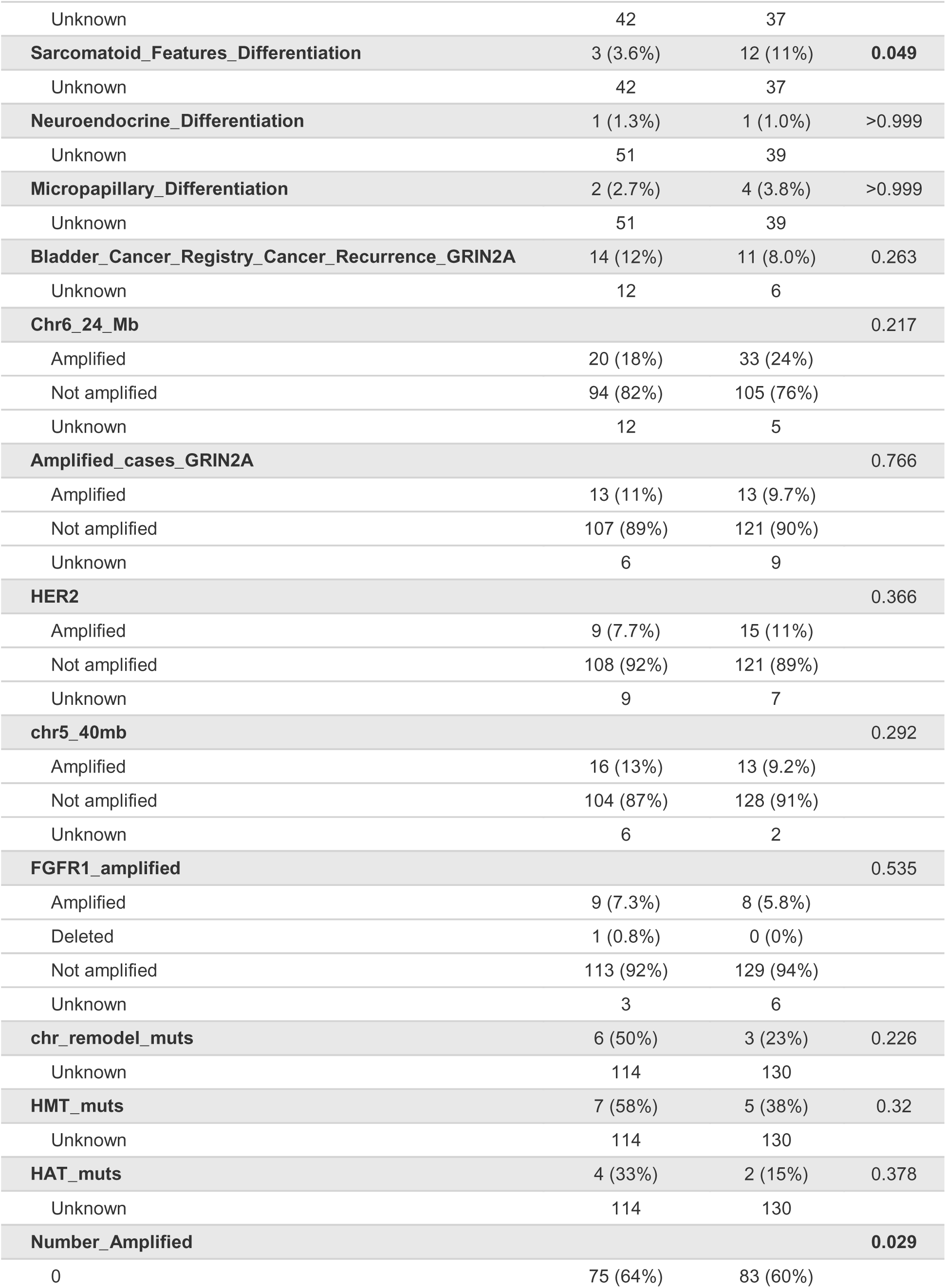
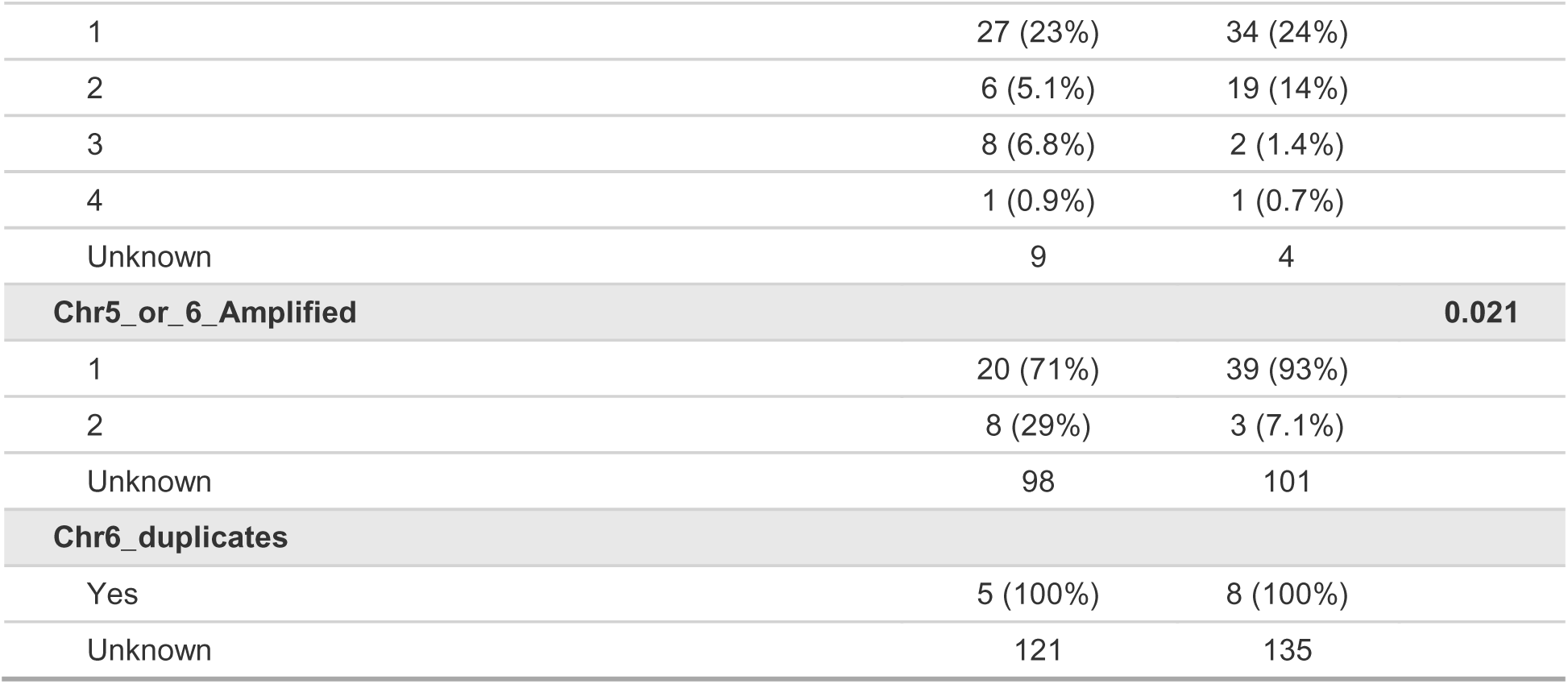
Statistical Comparisons of Patient and Tumor Characteristics based on SETDB1 H-Score of 10. Data presented as median (Q1, Q3) or n (%). P-values assessed via Wilcoxon rank sum test, Pearson’s Chi-squared test, or Fisher’s exact test where appropriate (n=269 patients).

SETDB1 classification was not significantly associated with tumor stage, suggesting that the altered survival outcomes in the SETDB1-high cohort are not due to a larger proportion of high-stage disease (Table 4). High SETDB1 expression was associated with increased frequency of squamous (35% vs 20%, p = 0.026) and sarcomatoid (11% vs 3.6%, p = 0.049) differentiation features (Table 4). Previous work has demonstrated that both squamous and sarcomatoid differentiation in bladder cancer is associated with higher risk of recurrence and poor survival [30, 31].

### SETDB1-high tumors are transcriptionally enriched for EMT, immune, and cisplatin resistance pathways

We found that SETDB1 is associated with shorter survival and altered differentiation, yet whether there were intrinsic differences in bladder tumors based on SETDB1 remained unknown. Therefore, we sought to determine SETDB1-specific tumor characteristics. To do this, we compared transcriptional profiles based on SETDB1 expression utilizing available RNA-sequencing data [32]. Of note, we did not detect a significant correlation between *SETDB1* mRNA expression and SETDB1 protein expression in our clinical samples (Figure S1E). Of the 269 patients with quantified SETDB1 H-scores, there were 34 cases with transcriptional data available (SETDB1 high: n = 29; SETDB1 low: n = 5). To identify SETDB1-associated transcriptional profiles, we performed differential expression analysis based on the H-score cutoff of 10 and identified differentially expressed genes (DEGs). SETDB1-high tumors had upregulation of 54 genes and downregulation of 20 genes (Figure S1F, Table S1). Gene set enrichment analysis showed that SETDB1-high tumors were enriched for the epithelial-to-mesenchymal transition pathway (NES = 1.95, p < 0.001), the interferon alpha response pathway (NES = 1.37, p = 0.018), and a curated gene signature [33] upregulated in the context of resistance to cisplatin treatment (NES = 1.88, p < 0.001) (Figure S1G). Given the small sample size, these data should be evaluated in a larger patient cohort for validation. Nonetheless, our results suggest that SETDB1 expression may be associated with altered transcriptional profiles in bladder tumors, which could potentially contribute to the altered survival we observe in our larger patient cohort.

## Discussion

Here, we find that high expression of SETDB1 in the tumors of bladder cancer patients is associated with significantly shorter survival, which held true upon multivariable analysis. Given the strong effects of SETDB1 on survival outcomes, this suggests that SETDB1 may be helpful as a prognostic biomarker for survival in bladder cancer, further supported by our ROC analyses in which SETDB1 performs modestly as a predictor of 2-, 3-, and 5-year patient survival. Bladder cancer is a molecularly heterogeneous disease, with widely varying mutational profiles and transcriptional states across patients [28, 34] and therefore identifying actionable biomarkers has proven difficult. The strong association we report here suggests that SETDB1 should be prospectively investigated as a protein biomarker in bladder cancer.

In addition to the possible predictive capabilities of evaluating SETDB1 expression in patient tumors, there is also precedent for targeting SETDB1 directly. In 2021, Guo et. al. described the first selective and potent SETDB1 inhibitor, SETDB1-TTD-IN-1 [20]. However, this inhibitor does not target the methyltransferase domain of SETDB1 and instead targets the tandem tudor domain responsible for histone binding. Since we do not yet know the precise mechanism by which SETDB1 is contributing to worse survival outcomes in bladder cancer, it is possible that this compound could be clinically relevant and therefore should be studied for efficacy in a preclinical setting. Further compound discovery studies should also be undertaken to detect SETDB1-specific methyltransferase inhibitors.

We found that higher SETDB1 expression is associated with increased frequency of aggressive bladder cancer subtypes, such as squamous and sarcomatoid, that are known to have altered transcriptional profiles [35]. Given that previous work has established the role of SETDB1 in maintaining stem cells via transcriptional regulation [9], we speculate that SETDB1 may be acting in a similar manner in bladder cancer to control differentiation state through regulating gene expression. Additionally, we assessed transcriptional differences between SETDB1-high and SETDB1-low tumors in a small cohort of samples that had available RNA-sequencing data, and identified enrichment of the EMT pathway, interferon alpha signaling, and genes associated with cisplatin resistance, further suggesting a role for SETDB1 in mediating gene expression in bladder cancer cells. SETDB1 has also been reported to methylate non-histone targets, such as p53 [36] and AKT [37], which could also contribute to an oncogenic function of SETDB1 in bladder cancer. Future studies should test whether altering SETDB1 expression affects differentiation states, transcriptional programs, and methylation of non-histone targets in bladder cancer.

A caveat of our study is retrospective design. Since all clinical information was collected at the time of diagnosis, some clinical information was unknown for individual patients, limiting the number of patients for group analyses. Additionally, due to the retrospective design the number of patients included in the transcriptional analysis of SETDB1-high versus SETDB1-low tumors were very small. Future studies should utilize a validation cohort with a representative number of both SETDB1-high and SETDB1-low samples to rigorously determine whether the transcriptional differences we observed in this small cohort will remain true in a larger sample size. To simplify associations with survival, we also computed and averaged scores for SETDB1 protein expression per patient which does not capture information on intra-tumor SETDB1 heterogeneity, which was visually observed in some cases. We did not observe a significant correlation between SETDB1 protein expression and *SETDB1* mRNA expression, which could be due to the small sample size or the use of average protein expression per patient. A larger cohort of patient samples with both protein- and RNA-level data, or evaluation of correlations within single-cell data would be required to assess whether there is a correlation between SETDB1 protein and RNA expression. Further, analysis of single-cell protein expression level would allow for analysis of intra-tumor heterogeneity of SETDB1 expression, which may reveal additional information as to how SETDB1 is mediating specific aspects of bladder tumor biology.

In conclusion, we found that higher tumor SETDB1 protein expression was independently associated with shorter overall survival in this retrospective bladder cancer cohort. Associations with squamous and sarcomatoid differentiation features and transcriptional programs warrant further investigation. Independent validation is needed to establish the prognostic value of SETDB1 protein expression.

## Acknowledgements

We acknowledge Dr. Wiam Bshara and Leighton Stein for their assistance in identifying regions of tumor on the evaluated TMAs and applying the Aperio algorithm for SETDB1 H-score quantification. This work was supported by National Cancer Institute (NCI) grants P30CA016056 and R25CA181003 involving the use of Roswell Park Comprehensive Cancer Center’s Biostatistics and Statistical Genomics and Biorepository and Laboratory Services Shared Resources.

---

**Figure S1. A.** Oncoprints for the TCGA Bladder Urothelial Carcinoma Firehose Legacy dataset (top) and MSK/TCGA Eur Urol 2020 dataset (bottom) showing the frequency of samples containing genomic alterations in SETDB1 and/or alterations in *SETDB1* mRNA expression. **B.** Representative images of tumor cores with high nuclear SETDB1 expression (left) and low nuclear SETDB1 expression (right), with the corresponding calculated H-score for the individual cores listed below. **C.** Log-rank statistics for log_2_(x+1)-transformed SETDB1 H-score, with the dotted line denoting the maximum statistic at 3.38, which is equivalent to a non-transformed H-score of 9.4. For simplicity, an H-score of 10 was utilized as the H-score cutoff for subsequent analyses. **D.** Receiver operating curve (ROC) for SETDB1 predictions of 2-, 3-, and 5-year survival. **E.** Scatter plot of SETDB1 H-score and normalized mRNA expression level of *SETDB1.* Pearson’s correlation coefficient and p-value displayed, computed using the stat_cor function from the ggpubr CRAN package in R. **F.** Volcano plot depicting differentially expressed genes (DEGs) between SETDB1-high (n=29) and SETDB1-low (n=5) tumors (full gene list in Table S1). Significantly upregulated genes (Log2FC > 1, padj < 0.05) are shown in red (n=54) and significantly downregulated genes (Log2FC < -1, padj < 0.05) are shown in blue (n = 20). For visualization efficiency, the highly significant gene *IRX1* (log_2_FC = 29.825, padj = 2.32e-13) is excluded from the plot. **G.** Gene set enrichment analysis plots for the Hallmark Epithelial to Mesenchymal Transition (left), Interferon Alpha Response (middle), and Kang Cisplatin Resistance Up (right) gene signatures.

**Table S1. SETDB1 High vs Low Differentially Expressed Genes (H-Score cut-off of 10).** Results of DESeq2 differential expression analysis RNA-seq data generated from patient tumors with high SETDB1 expression (H-score>10, n=29) or low SETDB1 expression (H score ≤10, n=5).

