## Supplementary figures and images for "Tumor SETDB1 expression predicts worse overall survival for bladder cancer patients"

### Supplemental Figure 1

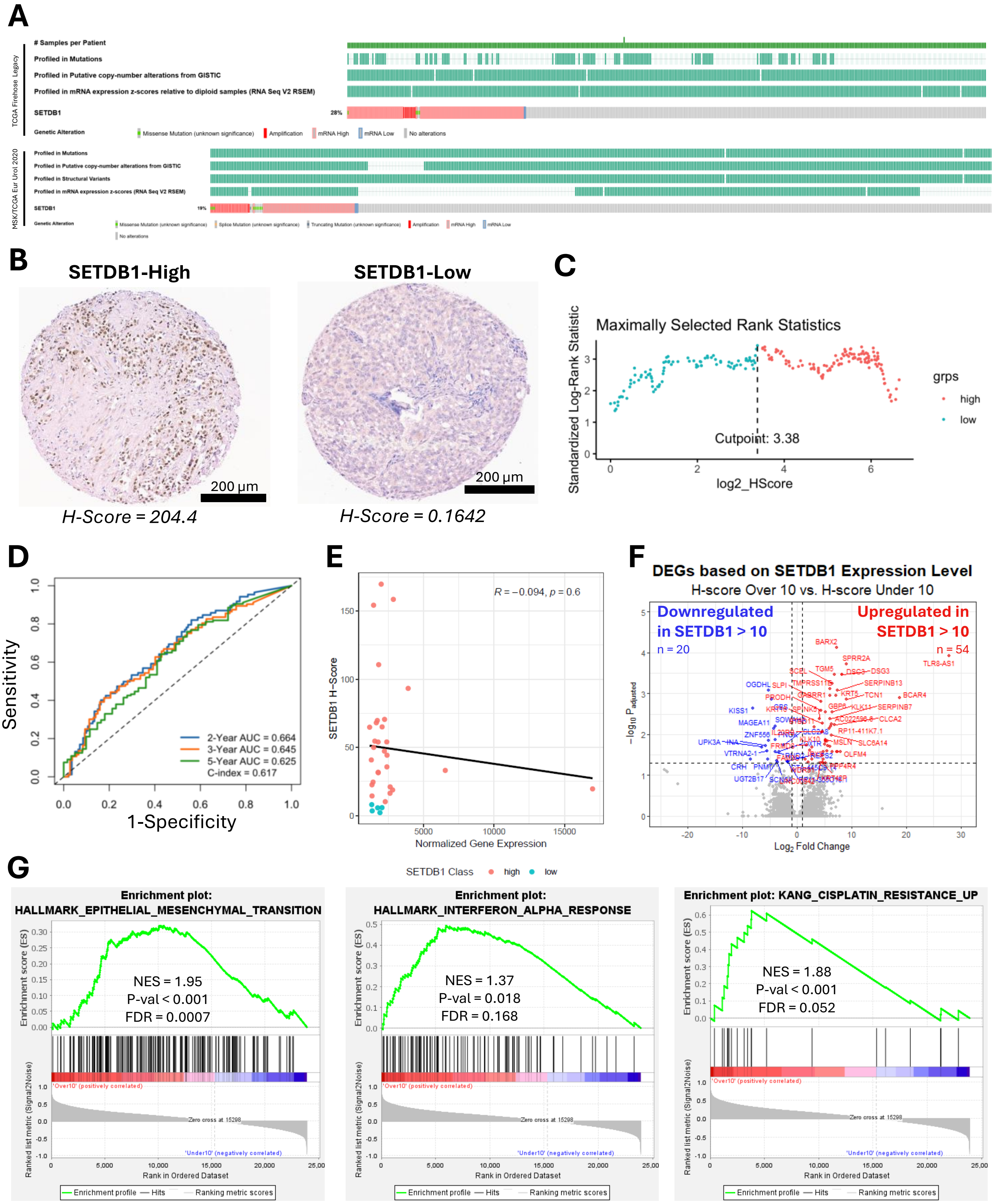
